# Lsm12 mediates Pol·q deubiquitination to help *Saccharomyces cerevisiae* resist oxidative stress

**DOI:** 10.1101/392415

**Authors:** Rui Yao, Liujia Shi, Chengjin Wu, Weihua Qiao, Liming Liu, Jing Wu

## Abstract

In *Saccharomyces cerevisiae*, the Y-family DNA polymerase η (Polη) regulates genome stability in response to different forms of environmental stress by translesion DNA synthesis. To elucidate the role of Polη in oxidative stress-induced DNA damage, we deleted or overexpressed the corresponding gene *RAD30*, and used transcriptome analysis to screen the potential genes associated with *RAD30* to respond to DNA damage. Under 2 mM H_2_O_2_, deletion of *RAD30* resulted in a 2.2-fold decrease in survival and a 2.8-fold increase in DNA damage, whereas overexpression of *RAD30* increased survival and decreased DNA damage by 1.2- and 1.4-fold, respectively, compared with that of the wild-type strain. Transcriptome and phenotypic analysis identified Lsm12 as a main factor involved in oxidative stress-induced DNA damage. Deleting *LSM12* caused growth defects while its overexpression enhanced cell growth under 2 mM H_2_O_2_. This effect was due to the physical interaction of Lsm12 with the UBZ domain of Polη to enhance Polη deubiquitination through Ubp3, and consequently promote Polη recruitment. Overall, these findings demonstrate that Lsm12 is a novel regulator mediating Polη deubiquitination to promote its recruitment under oxidative stress. Furthermore, this study provides a potential strategy to maintain the genome stability of industrial strains during fermentation.

**IMPORTANCE:** Polη was shown to be critical for cell growth in the yeast *Saccharomyces cerevisiae*, and deletion of its corresponding gene *RAD30* caused a severe growth defect under exposure to oxidative stress with 2 mM H_2_O_2_. Furthermore, we found that Lsm12 physically interacts with Polη and promotes Polη deubiquitination and recruitment. Overall, these findings indicate Lsm12 as a novel regulator mediating Polη deubiquitination that regulates its recruitment in response to DNA damage induced by oxidative stress.

## INTRODUCTION

Industrial microbial fermentation has been widely used in the production of chemicals. However, fermentation imposes a number of stresses on microorganisms, including oxidative stress, heat shock, osmotic stress, and exposure to toxic molecules and byproducts (1–3). Most of these factors form reactive oxygen species (ROS) that can cause DNA damage and genome instability, resulting in cell cycle arrest and cell death, thereby decreasing synthesis of the target compound (4,5). To solve this problem, cells have evolved a series of mechanisms for DNA damage tolerance.

In *Escherichia coli*, besides DNA repair mechanisms such as base excision repair and mismatch repair, there are two major pathways to deal with DNA damage: homology directed gap repair and translesion synthesis (TLS) (6). In the budding yeast *Saccharomyces cerevisiae*, there are three major strategies to maintain genome stability: template switch (TS) (7), homologous recombination (HR) (8), and TLS (9). TS is an error-free damage branch of the DNA damage tolerance mechanism, which is regulated by the polyubiquitination of proliferating cell nuclear antigen (PCNA) catalyzed by the Ubc13 and Mms2 enzymes (7,10,11). HR mainly repairs DNA double-strand breaks and is regulated by Srs2 and Rad51 (11). Srs2 is a DNA helicase that can bind with *SIZ1*-mediated sumoylated PCNA to prevent HR, and Rad51 is a recombinase that promotes HR (12). Similar to TS, HR also belongs to the error-free branch of the DNA damage tolerance pathway (13). In contrast, TLS is referred to as the error-prone branch of DNA damage tolerance (14), and is a conserved mechanism from bacteria to mammals that recruits various specialized DNA polymerases to the stalled replication forks (15–17). These specialized polymerases mostly belong to the Y family, consisting of Polη and Rev1 in yeasts, encoded by *RAD30* and *REV1*, respectively (18). The B family polymerase ξ, (Polξ is also involved in TLS (19).

Polη was first identified in yeast and has been shown to play a dominant role in DNA damage tolerance. Previous studies also demonstrated that Polη was particularly efficient at bypassing ultraviolet (UV) radiation-induced cyclobutane pyrimidine dimers, and could accurately insert an A opposite to the T of the dimer (20). Humans that lack Polη suffer from xeroderma pigmentosum variant, resulting in an extreme sensitivity to UV radiation (21). Polη can replicate 8-oxoguanine lesions efficiently and accurately by inserting a C opposite to the damage site (22). Polη can also bypass other lesions such as (6–4) TT photoproducts (23), O-6-methylguanine (24), abasic sites (25), and DNA double-strand breaks (26). In *S. cerevisiae*, Polη is recruited to stalled replication forks by its physical interaction with monoubiquitinated PCNA (27). However, the precise mechanism by which Polη is recruited to PCNA and its specific role in the response to oxidative stress-induced DNA damage is unclear. Therefore, in this study, we evaluated the role of Polη in H_2_O_2_-induced oxidative stress and analyzed the underlying mechanism.

## RESULTS

### *RAD30* is required for *S. cerevisiae* growth in the presence of H_2_O_2_

First, we checked whether *RAD30* is required for the growth of *S. cerevisiae* in the presence of H_2_O_2_. Toward this end, the wild-type, and *rad30Δ* and *rad30Δ/RAD30* mutant strains were spotted and grown on yeast nitrogen base medium with and without 2 mM H_2_O_2_ as a model of oxidative stress. Deletion of *RAD30* caused a significant growth defect in the presence of 2 mM H_2_O_2_, whereas overexpression of *RAD30* enhanced growth compared to that of the wild-type strain (Fig. 1A). Survival curves for all three strains were determined over a broad concentration range of H_2_O_2_ (Fig. 1B). At 2 mM H_2_O_2_, 70.4% of the wild-type strain survived, while the *rad30Δ* and *rad30Δ/RAD30* strains exhibited reduced (31.7%) and increased (84.5%) survival, representing a 2.2-fold decreased and 1.2-fold increase, respectively. These results suggest that *RAD30* contributes to cell growth in the presence of H_2_O_2_.

**Figure 1.**
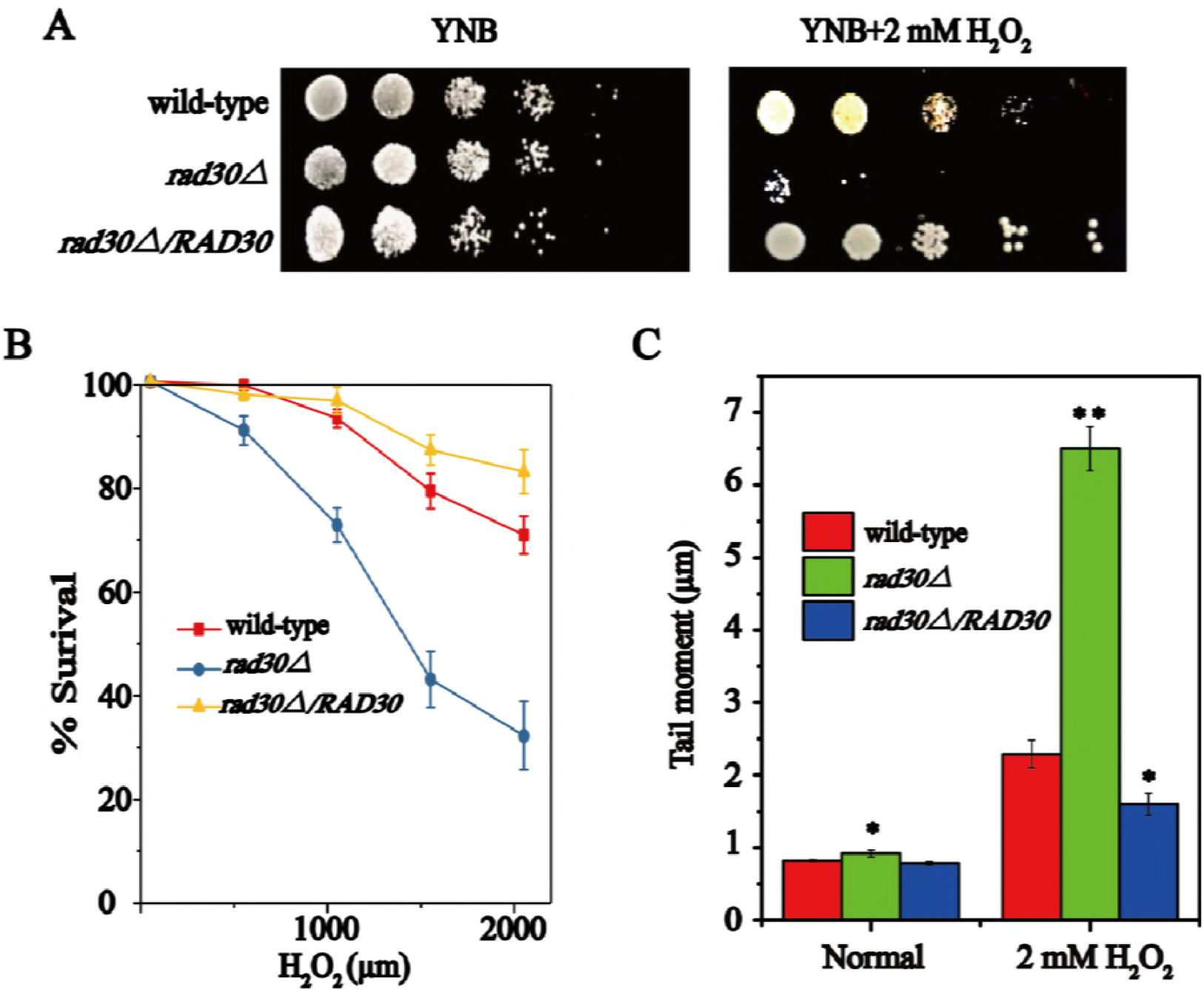
*RAD30* is required for *S. cerevisiae* growth in the presence of H_2_O_2_. (A) Wild-type, *rad30Δ*, and *rad30Δ/RAD30* strains were spotted on YNB plates under normal and 2 mM H_2_O_2_ conditions. (B) The survival rate of wild-type, *rad30Δ*, and *rad30Δ/RAD30* cells over a range of H_2_O_2_ doses (0, 500, 1000, 1500, 2000 μM). (C) Comet assay in wild-type, *rad30Δ*, and *rad30Δ/RAD30* strains exposed to normal or 2 mM H_2_O_2_ conditions. Data represent the means of three biological replicates (N = 3), and error bars represent SD. *P < 0.05, **P < 0.01.

To investigate the underlying mechanism, single-cell gel electrophoresis of the wild-type, *rad30Δ*, and *rad30Δ/RAD30* strains was performed. Without H_2_O_2_ treatment, both the *rad30Δ* and *rad30Δ/RAD30* strains displayed similar tail lengths relative to the wild-type strain. However, when treated with 2 mM H_2_O_2_, the *rad30Δ* and *rad30Δ/RAD30* strains showed a 2.8-fold increase and 1.4-fold decrease in tail length, respectively, when compared to that of the wild-type strain (Fig. 1C). This suggests that *RAD30* may play an important role in the response of *S. cerevisiae* to H_2_O_2_-induced DNA damage.

### Global transcriptome analysis of the *rad30Δ* and wild-type strains after treatment with H_2_O_2_

To further explain the weaker growth of the *rad30Δ* strain in the presence of H_2_O_2_, transcriptome sequencing was conducted to compare gene expression profiles in the *rad30Δ* and wild-type strains. We first compared the gene expression levels of wild-type cells at the log-phase of growth with and without H_2_O_2_ treatment. Transcriptional profiling analysis revealed 121 genes whose expression was significantly modified (|log2(fold change)| ≥ 1, FDR < 0.05), including 89 genes with up-regulated expression and 32 genes with down-regulated expression. In the *rad30Δ* strain, 804 genes displayed significantly differential expression, in which 424 were up-regulated and 380 were down-regulated (Fig. 2A). Specifically, there was a subset of 49 up-regulated and 15 down-regulated genes that were common to both the wild-type and *rad30Δ* strains (Table S1), indicating significant overlap. Gene Ontology (GO) analysis demonstrated that the commonly up-regulated genes were involved in DNA recombination process (GO:0006310), DNA damage response (GO:0006974), zinc ion homeostasis (GO:0006882), and oxidative stress response (GO:0006979), whereas the commonly down-regulated genes were enriched in GO processes such as cell wall chitin metabolism (GO:0006037), mitotic cell cycle (GO:0000278), and transport (GO: 0006810).

**Figure 2.**
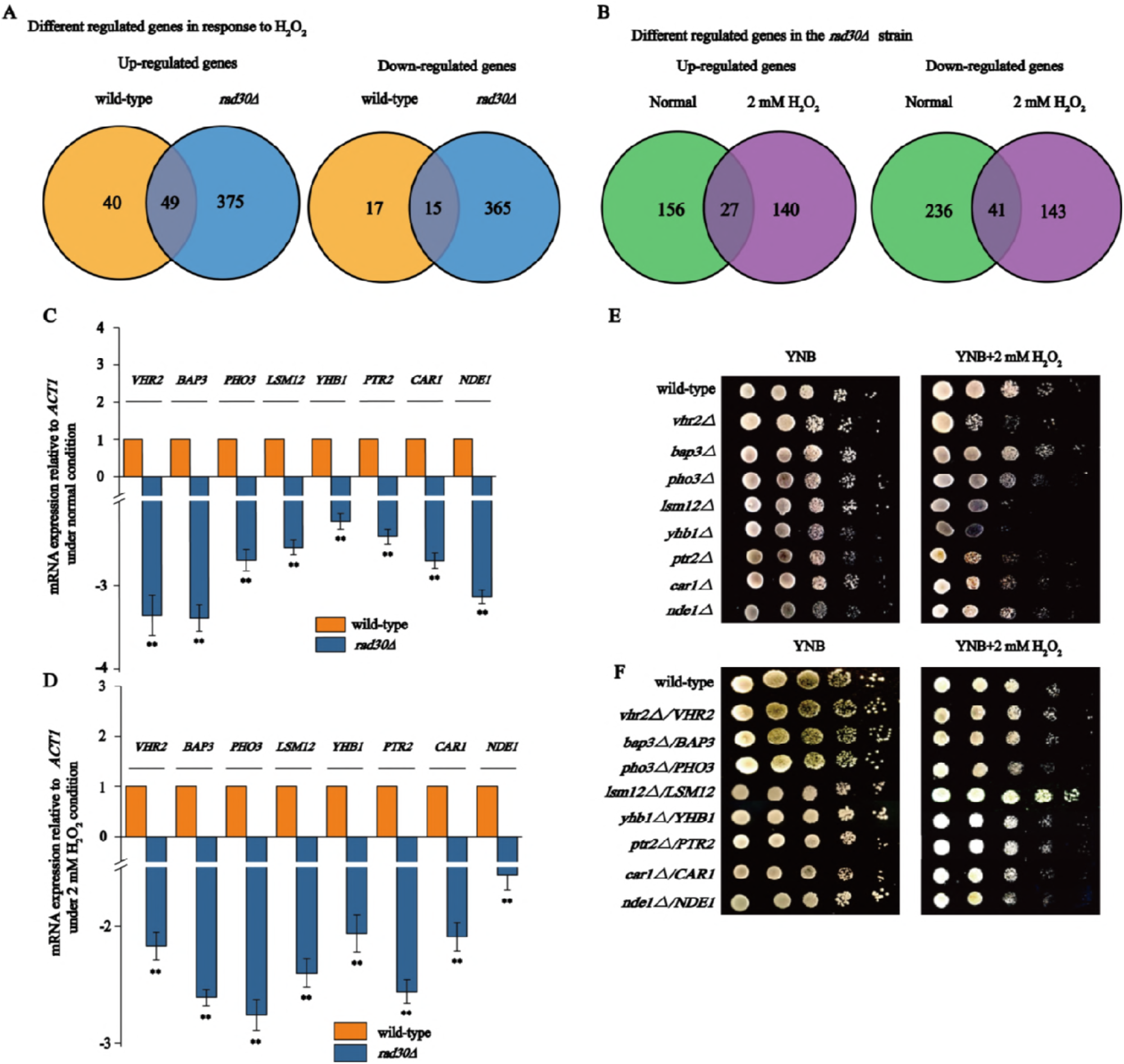
*LSM12* is involved in DNA damage tolerance. (A) Venn diagrams depicting the overlap between up-regulated and down-regulated genes in wild-type and *rad30Δ* strains in the normal condition compared with the gene expression levels in the corresponding strains at the 2 mM H_2_O_2_ condition. (B) Up-regulated and down-regulated genes in the *rad30Δ* mutant relative to their expression in the wild-type strain under normal and 2 mM H_2_O_2_ conditions. (C and D) qRT-PCR verified the mRNA expression levels of the most commonly down-regulated genes, calculated relative to the *ACT1* level, under normal and 2 mM H_2_O_2_ conditions. Data represent the means of three biological replicates (N = 3), and error bars represent SD. **P < 0.01. (E) The most commonly down-regulated genes were deleted and the mutant strains were spotted on YNB plates under normal and 2 mM H_2_O_2_ conditions. (F) The most commonly down-regulated genes were overexpressed and the mutant strains were spotted on YNB plates under normal and 2 mM H_2_O_2_ conditions.

Transcription profiling also revealed 183 and 277 genes with up-regulated and down-regulated expression, respectively, in the *rad30Δ* strain when compared to the wild-type strain without H_2_O_2_ treatment (Fig. 2B). However, when the cells were treated with 2 mM H_2_O_2_, 167 genes were up-regulated and 184 were down-regulated. There were subsets of 27 upregulated and 41 down-regulated genes that were common to both the normal and H_2_O_2_ conditions (Table S2), indicating significant overlap. GO analysis showed that the commonly up-regulated genes were involved in amino acid metabolism (GO:0006520), protein folding (GO:0006457), and DNA binding (GO:0003677) whereas the commonly down-regulated genes were enriched in processes such as meiosis I (GO:0007127), adenine metabolism (GO:0046083), DNA damage response (GO:0006974), and RNA metabolism (GO:0016070).

Among the genes commonly down-regulated in the *rad30Δ* strain, *VHR2, BAP3, PHO3, LSM12, YHB1, PTR2, CAR1*, and *NDE1* were the most significantly altered between the strains, with 3.36-, 3.42-, 2.84-, 2.56-, 2.22-, 2.49-, 2.77-, and 3.05-fold differences, respectively, under the normal condition, and with 2.2-, 2.63-, 2.79-, 2.41-, 1.85-, 2.55-, 2.09-, and 1.55-fold differences, respectively, under 2 mM H_2_O_2_. These results were further verified by reverse transcription-polymerase chain reaction (RT-PCR) analysis (Fig. 2C, D). To test whether these proteins interact with Polη or act in the same pathway, these genes were deleted or overexpressed in each strain, and the consequence on resistance to H_2_O_2_ stress was evaluated. Interestingly, deletion of *VHR2, LSM12*, or *YHB1* caused growth defects under the 2 mM H_2_O_2_ condition (Fig. 2E); however, only overexpression of *LSM12* conferred resistance to H_2_O_2_ (Fig. 2F). Based on these results, we hypothesized that *LSM12* may coordinate with *RAD30* to play an important role in DNA damage tolerance.

### Polη interacts with Lsm12 through the UBZ domain

On the basis of the above results, the subcellular localization of Polη and Lsm12 was determined. Under the normal condition, Lsm12 localized both in the nucleus and cytoplasm; however, following treatment with 2 mM H_2_O_2_, Lsm12 was mostly detected in the nucleus (Fig. 3A). In contrast, Polη was located in the nucleus both with and without H_2_O_2_ treatment. These results indicated that the relative distribution of Lsm12 in the nucleus increased with H_2_O_2_ treatment, supporting the hypothesis that Polη and Lsm12 may function together in the response to H_2_O_2_ treatment in the nucleus.

**Figure 3.**
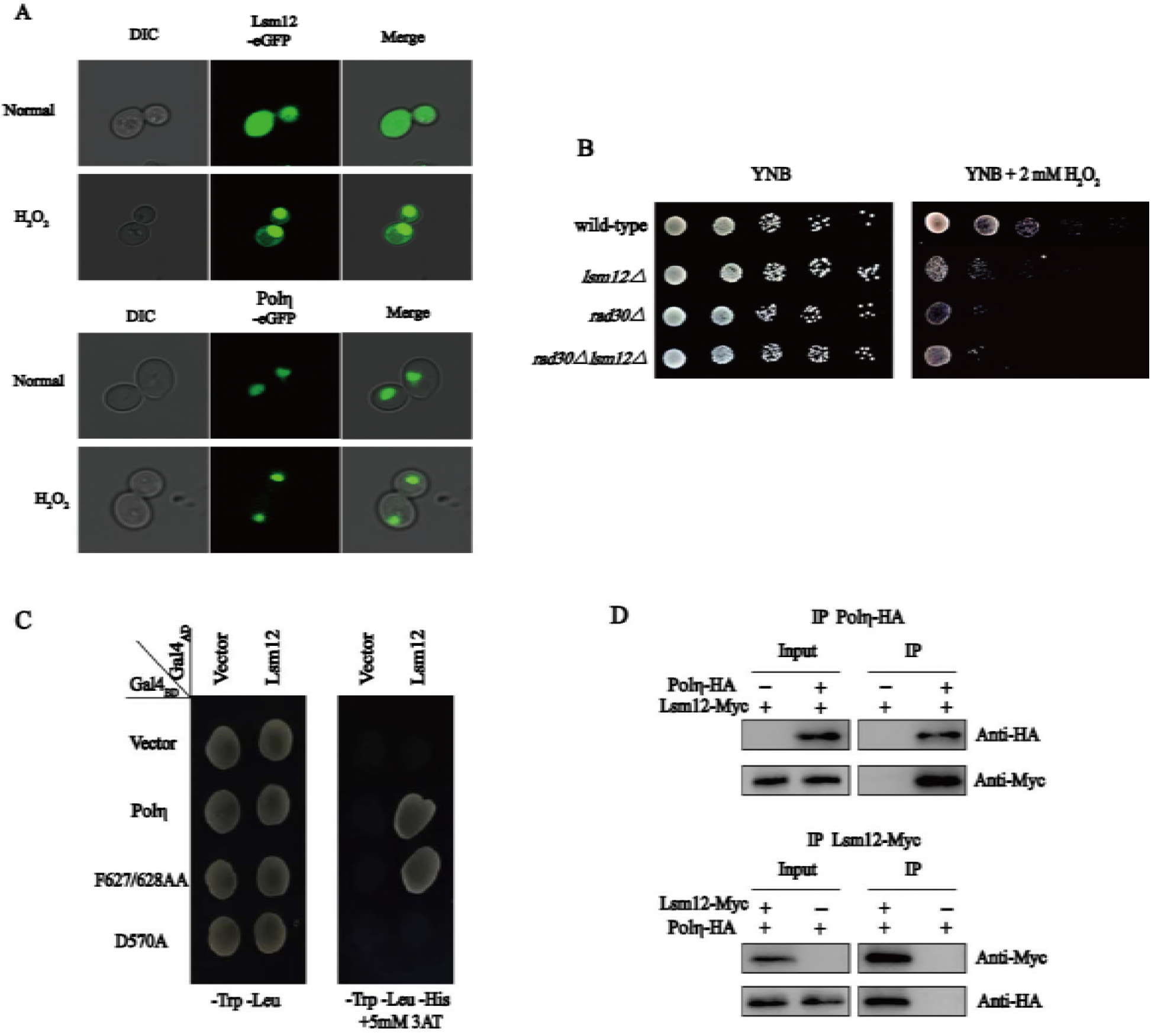
Polη interacts with Lsm12 through the UBZ domain. (A) Polη and Lsm12 were fused with the eGFP reporter and overexpressed, and the subcellular localization was visualized under normal and 2 mM H_2_O_2_ conditions. (B) The wild-type, *lsm12Δ*, *rad30Δ*, and *rad30Δlsm12Δ* strains were spotted on YNB plates with or without H_2_O_2_. (C) Yeast two-hybrid assays confirmed the interaction between Polη and Lsm12; the D570A mutant failed to interact with Lsm12. (D) Co-immunoprecipitation assay to detect the interaction between Polη and Lsm12 *in vivo.*

To further confirm this mechanism, we next examined the direct relationship between Lsm12 and Polη. First, the genetic interaction between Lsm12 and Polη was evaluated using spot assays, which revealed that the phenotype of the *rad30Δlsm12Δ* double mutant was similar to that of the *rad30Δ* and *lsm12Δ* single mutants (Fig. 3B). Moreover, the *rad30Δlsm12Δ* double mutant showed 33.6% survival, whereas the *rad30Δ* and *lsm12Δ* single mutants exhibited 31.7% and 36.5% survival, respectively (Table 1). These results demonstrated that the two genes have epistatic interactions.

**Table 1.** H_2_O_2_sensitivity of various yeast strains.

| Strain | Survival, % (without H<sub>2</sub>O<sub>2</sub>) | Survival, % (2 mM H<sub>2</sub>O<sub>2</sub>) |
| --- | --- | --- |
| wild-type | 100 | 70.4 (±3.9) |
| *rad30Δ* | 99.8(±1.64) | 31.7 (±4.9) ** |
| *rad30Δ/RAD30* | 98.1(±1.48) | 84.5 (±4.1) ** |
| *lsm12Δ* | 98.5(±2.26) | 36.5 (±4.7) ** |
| *rad30Δlsm12Δ* | 96.3(±0.75) | 33.6 (±2.2) ** |
| *ubp3Δ* | 98.1(±1.74) | 40.7 (±3.6) ** |
| *lsm12Δubp3Δ* | 96.1(±2.22) | 34.3 (±1.9) ** |

Survival rates, with the standard deviations shown, are expressed relative to those of wild-type
cells. Results are the average of three experiments. \*\* P values versus WT ≤ 0.01.

Next, the physical interaction between Lsm12 and Polη was determined. As shown in Fig. 3C, the yeast two-hybrid (Y2H) analysis revealed a gene-specific interaction between the full-length of Lsm12 and Polη. However, the D570A mutant, with an inactive UBZ domain of Polη, failed to interact with Lsm12. Furthermore, co-immunoprecipitation assays confirmed that Lsm12 and Polη physically interact *in vivo* (Fig. 3D), whereas this interaction did not occur with the D570A mutant, consistent with the Y2H results (data not shown). These observations suggest that Lsm12 physical interacts with Polη at the UBZ domain.

### Lsm12 promotes Polη recruitment in the presence of H_2_O_2_

Given the genetic and physical interaction between Lsm12 and Polη, we supposed that Lsm12 likely plays a role in DNA damage tolerance. Therefore, we next explored the mechanism by which Lsm12 repairs or facilitates tolerance to H_2_O_2_-induced DNA damage. Deletion of *LSM12* did not affect the mRNA or protein levels of Polη compared with those of the wild-type (data not shown). However, under H_2_O_2_ treatment, deletion of *LSM12* led to a decrease in the number of Polη foci formed with only 37.2%, in contrast to the 69.5% foci detected in the wild-type strain (Fig.4A and 4B). To further examine this result, the number of foci in the two strains after treatment with methyl methane sulfonate(MMS) were measured. Similarly, there were 76.2% and 43.3% Polη foci in the wild-type and *lsm12Δ* strain, respectively. These results suggest that Lsm12 promotes Polη recruitment to facilitate tolerance of DNA damage.

**Figure 4.**
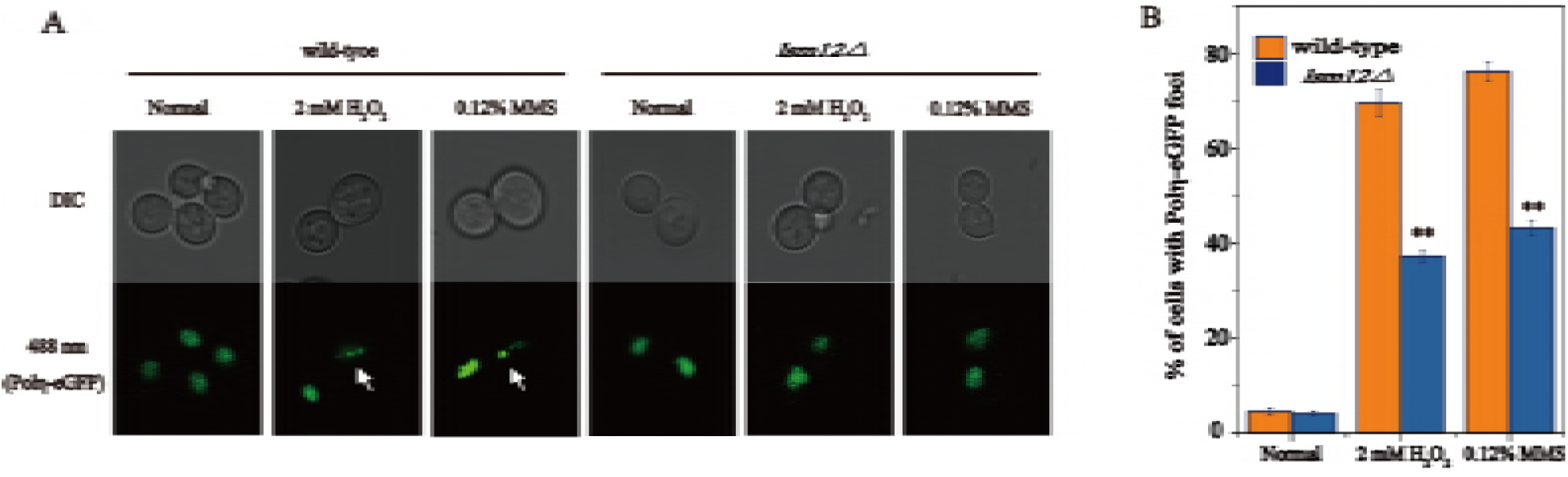
Lsm12 promotes Polη focus formation. (A) Formation of Polη foci when cells of wild-type and *lsm12Δ* strains were treated with different DNA-damaging agents. (B) Percentage of cells of different strains displaying Polη-eGFP foci in different environments. The histograms represent the mean ± SD from three independent experiments, **P < 0.01.

### Lsm12 deubiquitinates Polη through Ubp3

To elucidate the mechanism underlying the effect of Lsm12 in enhancing the formation of Polη foci in *S. cerevisiae*, the levels of PCNA and Polη monoubiquitination were compared in the wild-type and *lsm12Δ* strains without and with H_2_O_2_ treatment. As shown in Fig. 5A and 5B, the level of PCNA monoubiquitination significantly increased in both the wild-type (120%) and *lsm12Δ* (94%) strains after 2 mM H_2_O_2_ treatment, and there was no difference between the strains under either condition. By contrast, the level of Polη monoubiquitination significantly decreased in the wild-type (42%) after 2 mM H_2_O_2_ treatment, and was 102% higher in the *lsm12Δ* strain. This difference in the effects on PCNA and Polη monoubiquitination demonstrated that Lsm12 enhances Polη deubiquitination to promote Polη recruitment.

**Figure 5.**
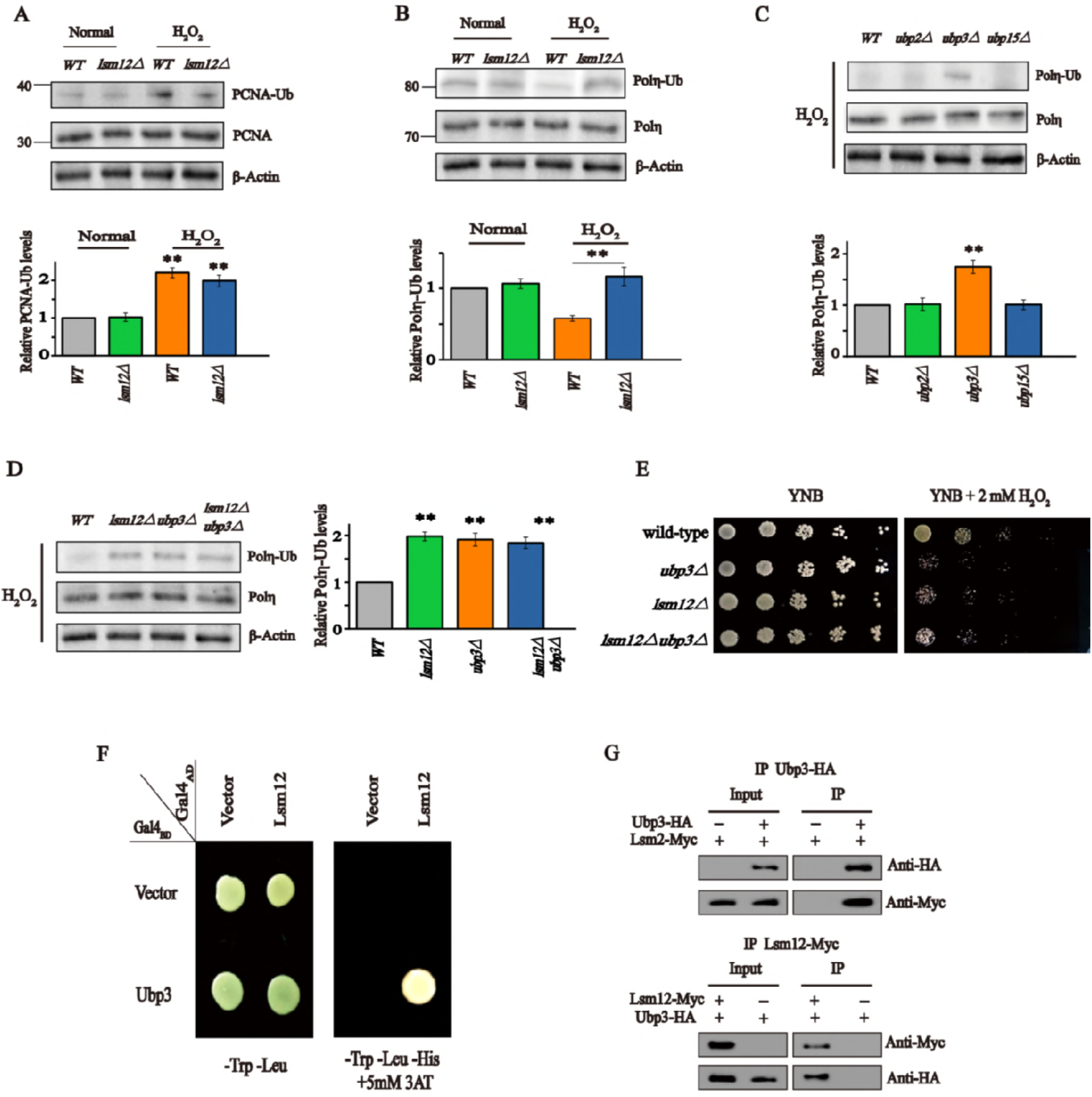
Lsm12 promoted Polη deubiquitination through Ubp3. (A) The level of monoubiquitinated PCNA in the wild-type and *lsm12Δ* strains. The level of monoubiquitinated Polη in the wild-type strain and (B) *lsm12Δ*; (C) *ubp2Δ, ubp3 Δ*, and *ubp15Δ*; and (D) *ubp3 Δ, lsm12Δ*, and *ubp3Alsm12Δ* strains. ß-Actin was used as a loading control. Data represent means ** of three biological replicates (N = 3), and error bars represent SD. P < 0.01. (E) Spot assays in 41 the wild-type, *ubp3Δ, lsm12Δ*, and *ubp3Δlsm12Δ* strains with or without H_2_O_2_. (F) Yeast two-hybrid assays confirmed the interaction between Lsm12 and Ubp3. (G) Co-immunoprecipitation assay to detect the interaction between Lsm12 and Ubp3 *in vivo.*

Given the lack of evidence that Lsm12 has its own deubiquitination activity, we hypothesized that Lsm12 binds with some deubiquitinase to catalyze the deubiquitination of Polη. To identify the specific deubiquitinase, we focused on the *UBP2, UBP3*, and *UBP15* genes, which are known to be associated with DNA damage tolerance. Under H_2_O_2_ treatment, deletion of *UBP2* and *UBP15* did not affect the level of Polη monoubiquitination compared with that of the wild-type, whereas deletion of *UBP3* increased Polη monoubiquitination (75%) (Fig. 5C).

Moreover, the level of Polη monoubiquitination in the *lsm12Δubp3Δ* double-mutant was similar to that of the *lsm12Δ* and *ubp3Δ* single-mutants under the H_2_O_2_ condition (Fig. 5D). Spot and survival assays also showed that the phenotype of the *lsm12Δubp3Δ* double-mutant was similar to that of the two single-mutants (Fig. 5E and Table 1). Further, both Y2H and co-immunoprecipitation experiments verified the physical interaction of Ubp3 with Lsm12 (Fig. 5F and 5G). These results suggest that Lsm12 promotes the deubiquitination of Polη, likely by binding with Ubp3.

## DISCUSSION

Translesion synthesis is a key pathway to maintain genome stability; however, the precise molecular mechanisms have not yet been clarified in detail. In this study, we demonstrated that deletion of *RAD30* caused a severe growth defect in the yeast *S. cerevisiae*, while its overexpression enhanced growth under oxidative stress due to exposure 2 mM H_2_O_2_. The stress response involves physical interaction between Lsm12 and Polη to tolerate or repair the consequent DNA damage. As a result, Lsm12 promoted Polη deubiquitination and facilitated Polη focus formation. These results demonstrate that Lsm12 mediates Polη deubiquitination and regulates its recruitment to help cells resist oxidative stress.

Previous studies have also indicated that *RAD30* appears to regulate cell growth under H_2_O_2_-induced DNA damage. In *S. cerevisiae*, cells lacking this gene are sensitive to UV radiation (28), MMS (29), and hydroxyurea (30). Yeast overexpressing Polη from *Trypanosoma cruzi* were reported to be more resistant to H_2_O_2_ exposure than the wild type (31). In human cells, loss of *POLH*, the orthologous gene to *RAD30* in *S. cerevisiae*, resulted in increased sensitivity to oxidative stress (32). Furthermore, knockdown of Polη in human cells decreased cell survival, and accelerated DNA damage and apoptosis (33). In our study, deletion of *RAD30* exhibited a severe growth defect, whereas overexpression of *RAD30* enhanced cell growth compared to that of the wild-type strain under 2 mM H_2_O_2_. This phenomenon was consistent with the previous findings in human cells, suggesting that Polη is a highly conserved protein from yeast to humans.

Lsm12 seems to be a multifunctional protein. Indeed, a previous study demonstrated that Lsm12 was involved in many aspects of RNA processing such as mRNA degradation, tRNA splicing, pre-mRNA splicing and degradation, and rRNA processing (34). In addition, Kim et al. (35) demonstrated that Lsm12 is involved in DNA replication stress. The present study provides new insight into this mechanism, showing that Lsm12 interacted with Polη to respond to the DNA damage induced by oxidative stress, and that this interaction occurs on the UBZ domain of Polη. In *S. cerevisiae*, Polη has two conserved domains, PIP and UBZ, encoded by the FF627, 628 and D570 residues, respectively (18). The PIP domain mainly interacts with monoubiquitinated PCNA when DNA is damaged (33). However, the function of the UBZ domain is not fully understood. A recent study showed that an inactive UBZ domain (*RAD30-D570A* mutant) failed to complement the phenotype of the *rad30Δ* mutant (36). Moreover, the UBZ domain of Polη was shown to be essential for 8-oxoguanine-induced mutagenesis (37).

Here, we demonstrated that Lsm12 promoted Polη deubiquitination and recruitment. When cells are under DNA replication stress, the Y family of DNA polymerases is recruited to the stalled replication forks (38). In this study, deletion of *LSM12* decreased the rate of Polη focus formation under the H_2_O_2_ condition, indicating that the absence of Lsm12 decreased Polη recruitment. This is like due to two mechanisms: (i) increasing PCNA monoubiquitination could promote Polη recruitment, because PCNA monoubiquitination can enhance affinity with Y family DNA polymerases (39), and Rad6/Rad18 induced PCNA monoubiquitination is essential for Polη recruitment (40); and (ii) decreasing Polη monoubiquitination could promote Polη recruitment. Previous studies indicated that when cells were exposed to UV radiation, the level of Polη monoubiquitination was down-regulated in the S-phase as a response to DNA damage (41). Similar results have also been detected in human cells (42). In this study, Lsm12 enhanced Polη recruitment through another mechanism given the observed decrease in the level of Polη monoubiquitination. However, this raises the question as to how Lsm12 deubiquitinates Polη. In *S. cerevisiae*, three deubiquitinases may be responsible for Polη deubiquitination: Ubp15, Ubp2, and Ubp3. Ubp15 leads to the accumulation of the mono-, di-, and poly-ubiquitination forms of PCNA (43). Ubp2 has been associated with oxidative stress, and the homologous gene in humans was shown to play a role in DNA damage tolerance (43). Ubp3 also appears to be involved in DNA replication stress given that a global protein abundance analysis revealed that the level of Ubp3 increased in response to exposure to DNA damage agents (44). Moreover, Ubp3 can stabilize Rad4 to enhance UV resistance, and promote the repair of UV-induced DNA damage (45). In this study, only the *ubp3Δ* mutant was found to increase the Polη monoubiquitination level, and genetic analyses further showed that *UBP3* and *LSM12* were epistatic. Accordingly, these two genes may function together in the deubiquitination of Polη. Both the Y2H and co-immunoprecipitation experiments confirmed a physical interaction between Lsm12 and Ubp3, which further validated our hypothesis.

In summary, we have identified a function of Lsm12 in the response to oxidative stress-induced DNA damage through interaction with Polη to promote Polη deubiquitination and recruitment. When cells were subjected to oxidative DNA replication stress, the amount of Lsm12 in the nucleus was increased, thereby promoting Polη deubiquitination and recruitment, to ultimately activate the TLS pathway and bypass DNA lesions. Cells with *LSM12* deleted failed to deubiquitinate Polη, leading to a defective TLS pathway (Fig. 6). These findings provide new insights into the molecular mechanisms of oxidative stress-induced DNA damage and suggest potential strategies to maintain the genomic stability of industrial strains.

**Figure 6.**
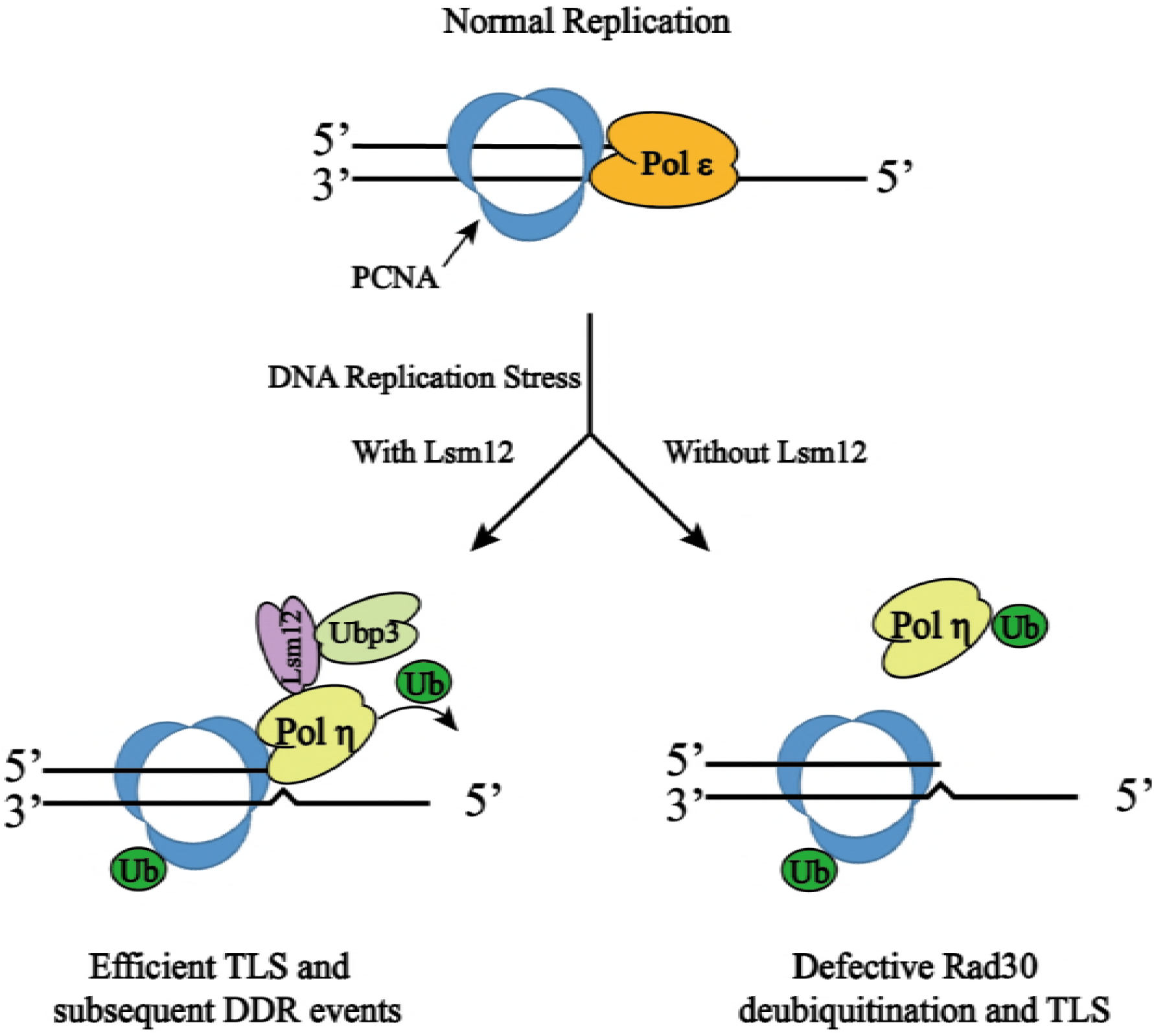
Model depicting the molecular function of Lsm12. When cells are under DNA replication stress, Lsm12 binds with Ubp3 and promote the deubiquitination of Polη, which activates the TLS pathway. In the absence of Lsm12, cells fail to deubiquitinate Polη, causing defective TLS.

## MATERIALS and METHODS

### Yeast strains, media and culture conditions

The *Saccharomyces cerevisiae* strains and plasmids used in this study are listed in Table S3. Null mutations were constructed by homologous recombination (46), and the overexpression constructs were made by either available restriction sites of predetermined fragments flanked by restriction sites in the primers. Site-specific mutations were made by the PCR-based method using the mutagenic primers. All primers used in this study are listed in Table S4 and S5.

Yeast cells were cultivated in Yeast extract peptone dextrose (YPD) medium (1% yeast extract, 2% Tryptone, 2% glucose, pH 6.5) and Yeast nitrogen base (YNB) medium (0.67% yeast nitrogen base without amino acid, 2% glucose, and supplemented with adenine (20.25 mg/liter), arginine (20 mg/liter), histidine (20 mg/liter), leucine (60 mg/liter), lysine (200 mg/liter), methionine (20 mg/liter), threonine (300 mg/liter), tryptophan (20 mg/liter), and uracil (20 mg/liter) pH 6.5). Yeast cells were grown at 30 °C with constant shaking at 200 rpm in a shaker-incubator chamber.

### Spot assays

Yeast cells cultivated in logarithmic phase, and diluted to an absorbance at 600 nm (A_600_) of 1.0 in phosphate-buffered saline (pH 7.0). Then, 10-fold serial dilutions cells were spotted onto YNB plates containing no drug or the indicated concentrations of H_2_O_2_. Growth was assessed after incubation for 2-4 days at 30 °C.

### Survival assays

Yeast cells cultivated in logarithmic phase, and harvested by centrifugation, washed with sterile water and resuspended in phosphate-buffered saline (pH 7.0) to 4 OD_600_ cells per milliliter. Cells were then treated with various doses of H_2_O_2_ for 1 h at 30 °C with 200 rpm shaking, followed by centrifugation and washing with sterile water for three times. After dilution, cells were plated in YNB medium plates and incubation at 30 °C for 2–4 days. Then the survival colonies were counted. Cells survival of each strain was expressed relative to that of untreated cells of the corresponding strain.

### Single cell gel electrophoresis

The Single cell gel electrophoresis was performed according to the protocol adopted for yeast cells (47). Approximately 10^6^ cells were harvested by centrifugation (2 min at 18000 g, 4 °C) and mixed with 1.5% (w/v) low melting agarose in S buffer (1 M sorbitol, 25 mM KH_2_PO_4_, pH 6.5) containing approximately 2 mg/mL of zymolyase (20T; 20000U/g). 80 μl of this mixture were spread over a slide coated with a water solution of 0.5% (w/v) normal melting agarose. Covered with a cover slip and incubated for 30 min at 30 °C for cell wall enzymatic degradation, after which the cover slips were removed. All further procedures were performed in a cold room at 4 °C. Slides were incubated in a lysis buffer (30 mM NaOH, 1 M NaCl, 0.05% laurylsarcosine, 50 mM EDTA, 10 mM Tris-HCl, pH 10) for 2 h in order to lyse yeast spheroplasts. The slides were washed three times for 20 min each in electrophoresis buffer (30 mM NaOH, 10 mM EDTA, 10 mM Tris-HCl, pH 10) to remove lysis solution. The slides were then submitted to electrophoresis in the same buffer for 15 min at 0.7 V/cm. After electrophoresis, the slides were incubated in a neutralization buffer (10 mM Tris-HCl, pH 7.4) for 10 min, followed by consecutive 5 min incubation in 76% and 96% ethanol. The slides were then air-dried and were visualized immediately or stored at 4 °C for later observation. For visualization in a fluorescence microscope the slides were stained with ethidium bromide (10 μg/mL) and 20 representative images of each slide were acquired at magnification of 400× using a Leica Microsystems DM fluorescence microscope. The images were analyzed with the help of the free edition of Comet Assay Software Project (CASP) and the analytic parameter Tail Length (in μm) was chosen as the unit of DNA damage. In each slide, at least 20 comets were analyzed and error bars represent variability between the mean of at least three different slides obtained from biologically independent experiments.

### Genome-wide transcription analysis

The wide-type, *rad30Δ* strains cultivated in logarithmic phase, and then H_2_O_2_ was added for a final concentration of 2 mM. Cells collected after 1 h of H_2_O_2_ treatment. Total RNA was isolated by using MiniBEST Universal RNA Extraction Kit (TaKaRa Bio, Shiga, Japan). The concentration and quality of total RNA were determined by microspectrophotometry using an Agilent 2100 Bioanalyzer (Agilent Technologies, Santa Clara, CA). Frozen samples were sent to the Majorbio Institute (http://www.majorbio.com/) for global gene analysis. The raw date is available at https://www.ncbi.nlm.nih.gov/sra/SRP151558, detailed descriptions are included in supplementary materials. The annotation and the Gene Ontology (GO) were based on the *Saccharomyces* Genome Database (SGD).

### qRT-PCR analysis

Cells cultivated in logarithmic phase, and treated with 2 mM H_2_O_2_ for 1 h. Total RNA was extracted by using the MiniBEST Universal RNA Extraction Kit (TaKaRa Bio, Shiga, Japan). And the cDNA was synthesized by using the PrimeScript II 1st-strand cDNA synthesis kit (6210A; TaKaRa Bio). The quantitation of mRNA level was analyzed by using SYBR Premix ExTaq (RR420A; Takara Bio). *ACT1* as a standard control to normalized the gene expression.

### Yeast two-hybrid (Y2H) assays

All Y2H plasmids were based on either *pGBKT7* (Gal4_BD_) or *pGADT7* (Gal4_AD_). The *pGBKT7-RAD30, pGADT7-LSM12, pGADT7-UBP3*, and other point mutant fusion proteins plasmids were constructed by standard genetic techniques. Gal4_AD_ and Gal4_BD_ plasmids to be tested were co-transformed into the yeast strain *AH109*, individual colonies were picked and then allowed to grow at 30 °C on an SD-Leu-Trp plate for 2–3 days, after which transformants were printed on SD-Leu-Trp, SD-Leu-Trp-Ade and SD-Leu-Trp-His selective plates with or without a certain amount of the histidine biosynthesis inhibitor 1,2,4-aminotrizole (3-AT) (48).

### Western blotting

Polη and PCNA containing C-terminal HA tag were expressed from its native promoter in wild-type and *lsm12Δ* strains. Cells were grown to logarithmic phase and harvested by centrifugation, than resuspended in lysis buffer containing 50 mM TrisHCl (pH 7.5), 150 mM NaCl, 1 mM EDTA, 10% (vol/vol) glycerol, 0.1% Tween 20, 1 mM phenylmethylsulfonyl fluoride (PMSF), and 1 × complete protease inhibitor mixture (Sangon Biotech). Cells were broken by bead beating (45 min at 4 °C) with glass beads and collected the supernatant. The extracts were resolved by SDS-PAGE in 10% acrylamide gels, and transferred to a PVDF membrane, blocked with 5% milk in TBST. The monoubiqination level of Polη and PCNA were probed with mouse Anti-HA tag antibody (ab18181) and rabbit Anti-Mouse IgG secondary antibodies conjugated HRP (ab6728). The bands were visualized using a ChemiDoc™ XRS+ imaging system.

### Co-immunoprecipitation

Cells transformed with indicated plasmids and total proteins were extracted by lysis buffer. The extracts were incubated with 25 μL anti-HA-conjugated magnetic beads (Bio-Rad) over night at 4 °C and washed three times by lysis buffer. Next the precipitates were eluted into 100 mM Glycine (pH 2.5) and 100 mM NaCl, immediately neutralized by 2 M TrisHCl (pH 9.0) and 100 mM NaCl, and finally performed the immunoblot analysis.

### Microscope analysis

The method followed as described previously (49, 50) with slight modifications. Yeast strains cultivated in logarithmic phase, incubation with 2 mM H_2_O_2_ for 2 h or 0.12% MMS for 1 h. Then cells were harvested by centrifuging and washed with 0.1 M phosphate buffer (PBS, pH 7.5). The pellet was resuspended in 20 μL 0.1 M PBS with 1.2 M sorbitol for microscopy observation. Images were obtained with Leica TCS SP8 confocal microscope, using 488 nm for eGFP. The percentage of cells with foci was counted in three independent experiments, and at least 500 cells per experiment.

### Quantification and statistical analysis

For quantification of the western blot data, Image J software was used to measure the relative intensity of each band, and the relative PCNA-Ub and Polη-Ub protein levels were normalized to the relative ß-Actin levels. Quantification data were presented as the mean ± SD (standard deviation) from at least three independent experiments. Statistical differences were determined by the *t* test.

## ACKNOWLEDGMENTS

This work was supported by the National Natural Science Foundation of China (21706095, 21676118), Jiangsu Province “333 High-level Talents Cultivating Project” (BRA2016365), and the national first-class discipline program of Light Industry Technology and Engineering (LITE2018-20).

## Author contributions

R.Y., L.L. and J.W. designed research; R.Y., L.S., C.W., and W.Q. performed research; R.Y. and J.W. analyzed data; and R.Y. and L.L. wrote the paper.

---

**Table S3.**
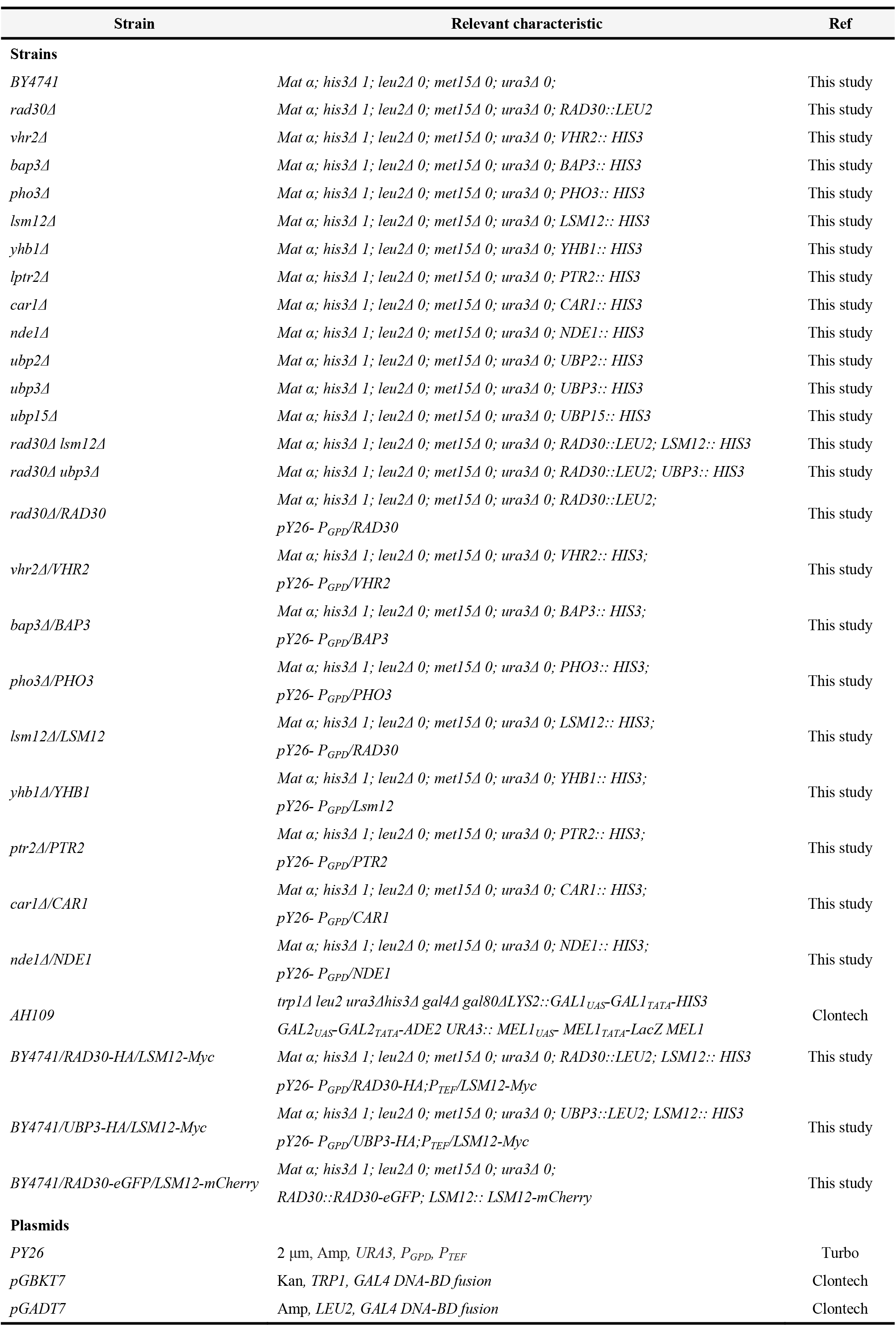
Strains and Plasmids ids used in this study.

**Table S4.**
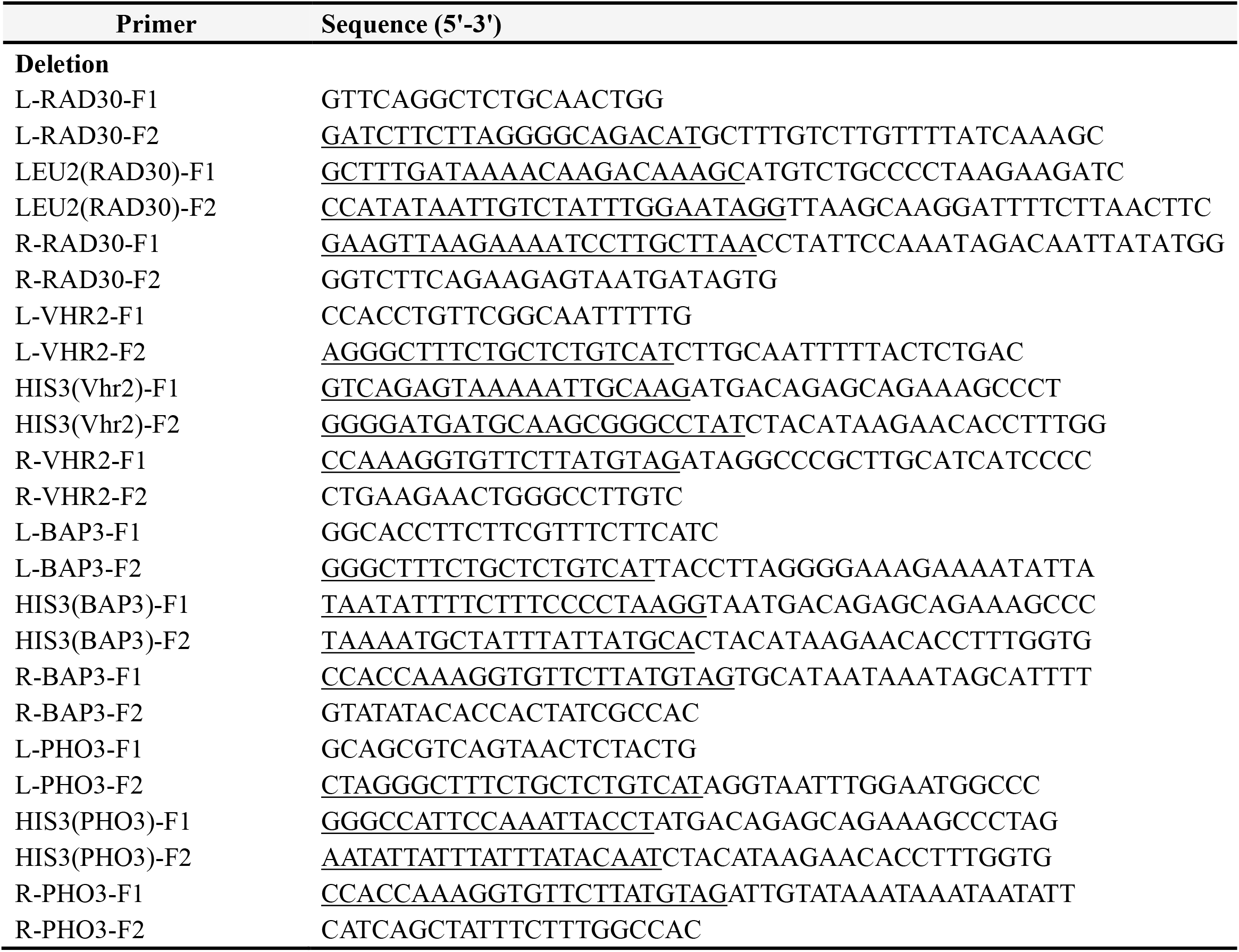
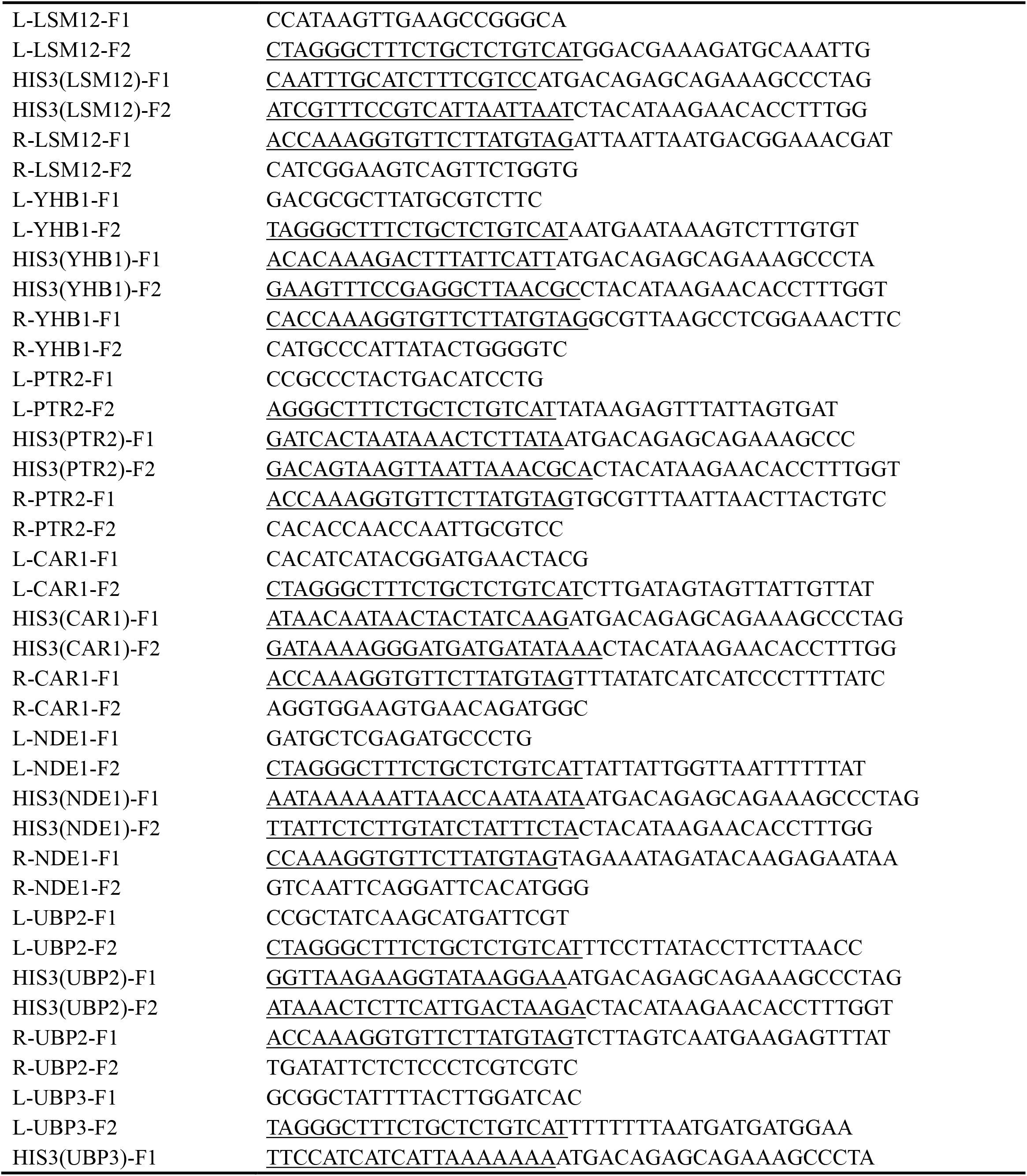
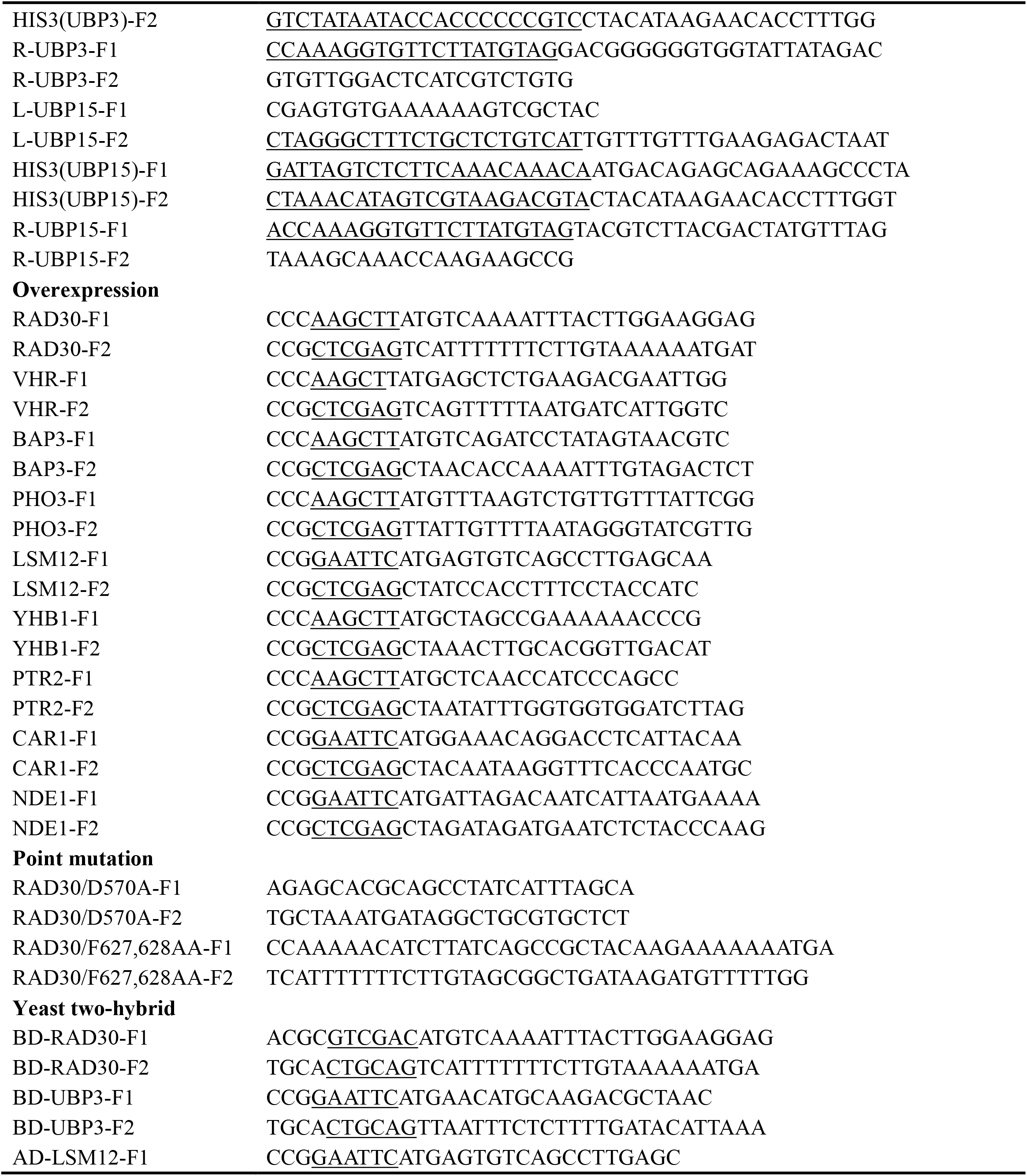
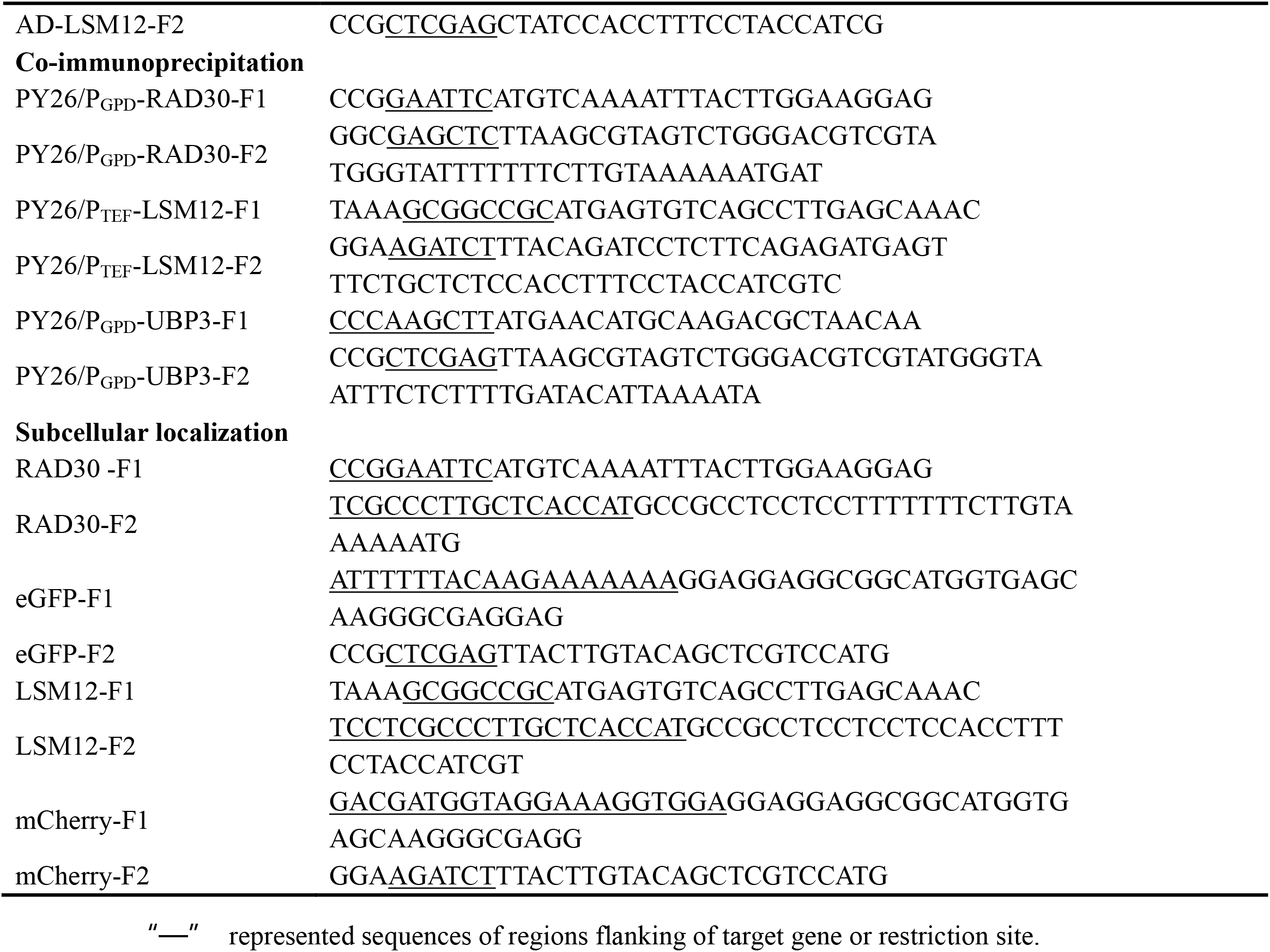
Primers used for plasmid construction in this study.

**Table S5.**
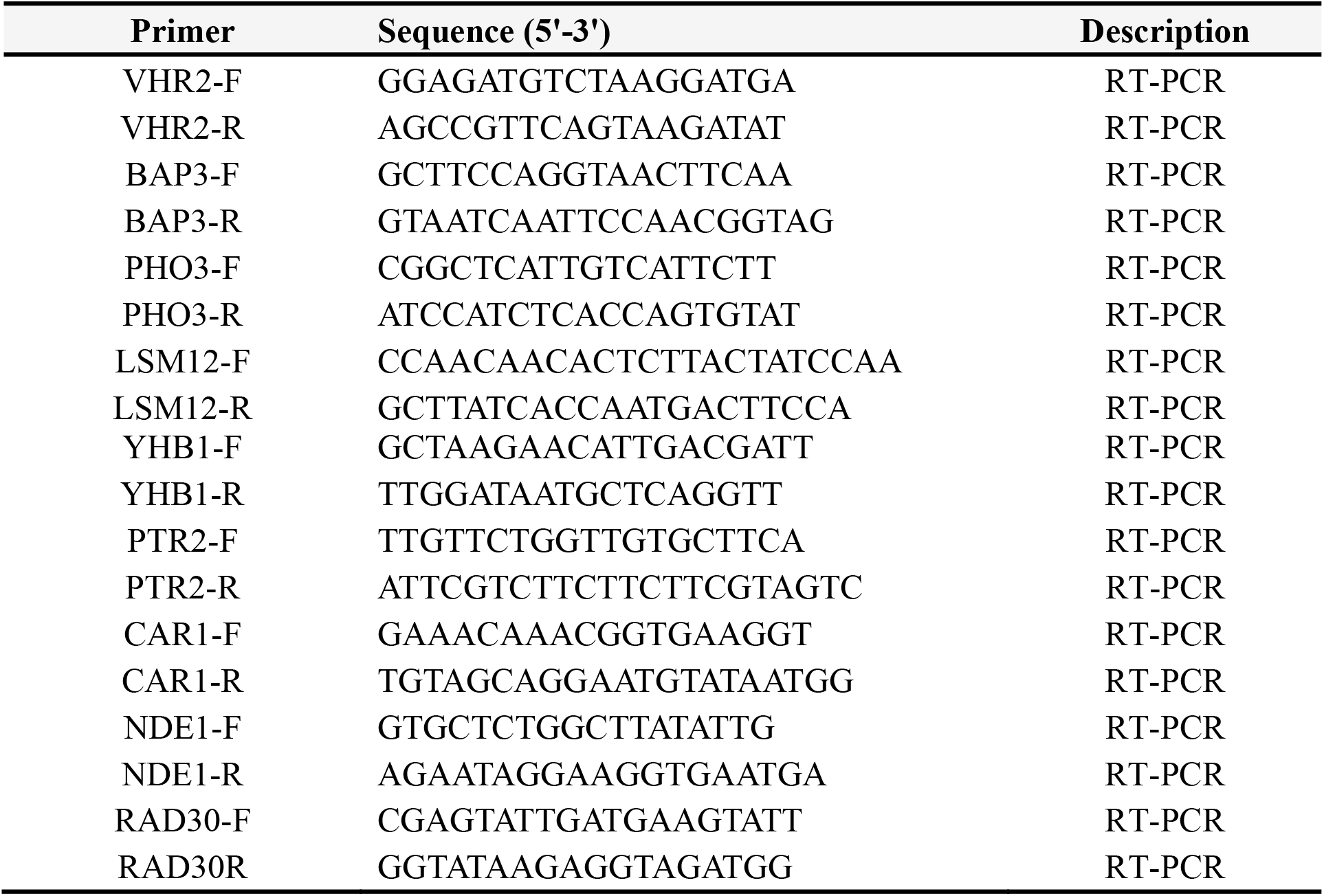
Primers used for RT-PCR in this study

## REFERENCES

1. Novarina D, Mavrova SN, Janssens GE, Rempel IL, Veenhoff LM, Chang M. 2017. Increased genome instability is not accompanied by sensitivity to DNA damaging agents in aged yeast cells. DNA Repair 54:1–7.

2. Hayashi MT, Cesare AJ, Rivera T, Karlseder J. 2015. Cell death during crisis is mediated by mitotic telomere deprotection. Nature 522:492–496.

3. Saini P, Beniwal A, Vij S. 2017. Physiological response of Kluyveromyces marxianus during oxidative and osmotic stress. Process Biochem 56:21–29.

4. Zhao HW, Li JY, Han BZ, Li X, Chen JY. 2014. Improvement of oxidative stress tolerance in Saccharomyces cerevisiae through global transcription machinery engineering. J Ind Microbiol Biot 41:869–878.

5. Roukas T. 2016. The role of oxidative stress on carotene production by Blakeslea trispora in submerged fermentation. Criti Rev Biotechno 36:424–433.

6. Naiman K, Pages V, Fuchs RP. 2016. A defect in homologous recombination leads to increased translesion synthesis in E. coli. Nucleic Acids Res 44:7691–7699.

7. Fumasoni M, Zwicky K, Vanoli F, Lopes M, Branzei D. 2015. Error-free DNA damage tolerance and sister chromatid proximity during DNA replication rely on the pol alpha/Primase/Ctf4 complex. Mol Cell 57:812–823.

8. Villoria MT, Ramos F, Duenas E, Faull P, Cutillas PR, Clemente-Blanco A. 2017. Stabilization of the metaphase spindle by Cdc14 is required for recombinational DNA repair. EMBO J 36:79–101.

9. Shemesh K, Sebesta M, Pacesa M, Sau S, Bronstein A, Parnas O, Liefshitz B, Venclovas C, Krejci L, Kupiec M. 2017. A structure-function analysis of the yeast Elg1 protein reveals the importance of PCNA unloading in genome stability maintenance. Nucleic Acids Res 45:3189–3203.

10. Gervai JZ, Galicza J, Szeltner Z, Zamborszky J, Szuts D. 2017. A genetic study based on PCNA-ubiquitin fusions reveals no requirement for PCNA polyubiquitylation in DNA damage tolerance. DNA Repair 54:46–54.

11. Northam MR, Trujillo KM. 2016. Histone H2B mono-ubiquitylation maintains genomic integrity at stalled replication forks. Nucleic Acids Res 44:9245–9255.

12. Zhang SC, Wang LL, Tao Y, Bai TY, Lu R, Zhang TL, Chen JY, Ding JP. 2017. Structural basis for the functional role of the Shu complex in homologous recombination. Nucleic Acids Res 45: 13068–13079.

13. Budzowska M, Graham TGW, Sobeck A, Waga S, Walter JC. 2015. Regulation of the Rev1-pol zeta complex during bypass of a DNA interstrand cross-link. EMBO J 34:1971–1985.

14. Rechkoblit O, Kolbanovskiy A, Landes H, Geacintov NE, Aggarwal AK. 2017. Mechanism of error-free replication across benzo [a] pyrene stereoisomers by Rev1 DNA polymerase. Nat Commun 8:965.

15. Huang M, Zhou B, Gong J, Xing L, Ma X, Wang F, Wu W, Shen H, Sun C, Zhu X, Yang Y, Sun Y, Liu Y,Tang T-S, Guo C. 2018. RNA-splicing factor SART3 regulates translesion DNA synthesis. Nucleic Acids Res 46:4560–4574.

16. Yang Y, Gao Y, Mutter-Rottmayer L, Zlatanou A, Durando M, Ding W, Wyatt D, Ramsden D, Tanoue Y, Tateishi S, Vaziri C. 2017. DNA repair factor Rad18 and DNA polymerase pol kappa confer tolerance of oncogenic DNA replication stress. J Cell Biol 216:3097–3115.

17. Vujanovic M, Krietsch J, Raso MC, Terraneo N, Zellweger R, Schmid JA, Taglialatela A, Huang J-W, Holland CL, Zwicky K, Herrador R, Jacobs H, Cortez D, Ciccia A, Penengo L, Lopes M. 2017. Replication fork slowing and reversal upon DNA damage require PCNA polyubiquitination and ZRANB3 DNA translocase activity. Mol Cell 67:882–890.

18. Vaisman A, Woodgate R. 2017. Translesion DNA polymerases in eukaryotes: what makes them tick? Crit Rev Biochem Mol 52:274–303.

19. Tellier-Lebegue C, Dizet E, Ma E, Veaute X, Coic E, Charbonnier JB, Maloisel L. 2017. The translesion DNA polymerases pol zeta and Rev1 are activated independently of PCNA ubiquitination upon UV radiation in mutants of DNA polymerase delta. Plos Genet 13:e1007119.

20. Sale JE, Lehmann AR, Woodgate R. 2012. Y-family DNA polymerases and their role in tolerance of cellular DNA damage. Nat Rev Mol Cell Bio 13:141–152.

21. De Palma A, Morren MA, Ged C, Pouvelle C, Taieb A, Aoufouchi S, Sarasin A. 2017. Diagnosis of xeroderma pigmentosum variant in a young patient with two novel mutations in the POLH gene. Am J Med Genet A 173:2511–2516.

22. Xue QZ, Zhong MY, Liu BY, Tang Y, Wei ZL, Guengerich FP, Zhang HD. 2016. Kinetic analysis of bypass of 7, 8-dihydro-8-oxo-2’-deoxyguanosine by the catalytic core of yeast DNA polymerase eta. Biochimie 121:161–169.

23. Choi JH, Pfeifer GP. 2005. The role of DNA polymerase eta in UV mutational spectra. DNA Repair 4:211–220.

24. Liu BY, Xue QZ, Gu SL, Wang WP, Chen J, Li YQ, Wang CX, Zhang HD. 2016. Kinetic analysis of bypass of O-6-methylguanine by the catalytic core of yeast DNA polymerase eta. Arch Biochem Biophys 596:99–107.

25. Yang JT, Wang R, Liu BY, Xue QZ, Zhong MY, Zeng H, Zhang HD. 2015. Kinetic analysis of bypass of abasic site by the catalytic core of yeast DNA polymerase eta. Mutat Res-Fund Mol M 779:134–143.

26. Saito Y, Zhou H, Kobayashi J. 2015. Chromatin modification and NBS1: their relationship in DNA double-strand break repair. Genes Genet Syst 90:195–208.

27. Boehm EM, Powers KT, Kondratick CM, Spies M, Houtman JCD, Washington MT. 2016. The proliferating cell nuclear antigen (PCNA)-interacting protein (PIP) motif of DNA polymerase mediates its interaction with the C-terminal domain of Rev1. J Biol Chem 291:8735–8744.

28. Donigan KA, Cerritelli SM, McDonald JP, Vaisman A, Crouch RJ, Woodgate R. 2015. Unlocking the steric gate of DNA polymerase eta leads to increased genomic instability in Saccharomyces cerevisiae. DNA Repair 35:1–12.

29. Wit N, Buoninfante OA, van den Berk PCM, Jansen JG, Hogenbirk MA, de Wind N, Jacobs H. 2015. Roles of PCNA ubiquitination and TLS polymerases kappa and eta in the bypass of methyl methanesulfonate-induced DNA damage. Nucleic Acids Res 43:282–294.

30. Chen X, Bosques L, Sung P, Kupfer GM. 2016. A novel role for non-ubiquitinated FANCD2 in response to hydroxyurea-induced DNA damage. Oncogene 35:22–34.

31. de Moura MB, Fonseca Schamber-Reis BL, Passos Silva DG, Rajao MA, Macedo AM, Franco GR, Junho Pena SD, Ribeiro Teixeira SM, Machado CR. 2009. Cloning and characterization of DNA polymerase eta from Trypanosoma cruzi: roles for translesion bypass of oxidative damage. Environ Mol Mutagen 50:375–386.

32. Zlatanou A, Despras E, Braz-Petta T, Boubakour-Azzouz I, Pouvelle C, Stewart GS, Nakajima S, Yasui A, Ishchenko AA, Kannouche PL. 2011. The hMsh2-hMsh6 complex acts in concert with monoubiquitinated PCNA and Pol eta in response to oxidative DNA damage in human cells.Mol Cell 43:649–662.

33. Boehm EM, Spies M, Washington MT. 2016. PCNA tool belts and polymerase bridges form during translesion synthesis. Nucleic Acids Res 44:8250–8260.

34. Fleischer TC, Weaver CM, McAfee KJ, Jennings JL, Link AJ. 2006. Systematic identification and functional screens of uncharacterized proteins associated with eukaryotic ribosomal complexes. Gene Dev 20:1294–1307.

35. Lee MW, Kim BJ, Choi HK, Ryu MJ, Kim SB, Kang KM, Cho EJ, Youn HD, Huh WK, Kim ST. 2007. Global protein expression profiling of budding yeast in response to DNA damage. Yeast 24:145–154

36. Acharya N, Brahma A, Haracska L, Prakash L, Prakash S. 2007. Mutations in the ubiquitin binding UBZ motif of DNA polymerase eta do not impair its function in translesion synthesis during replication. Mol Cell Biol 27:7266–7272.

37. van der Kemp PA, de Padula M, Burguiere-Slezak G, Ulrich HD, Boiteux S. 2009. PCNA monoubiquitylation and DNA polymerase eta ubiquitin-binding domain are required to prevent 8-oxoguanine-induced mutagenesis in Saccharomyces cerevisiae. Nucleic Acids Res 37:2549–2559.

38. Enervald E, Lindgren E, Katou Y, Shirahige K, Strom L. 2013. Importance of Pol eta for damage-induced cohesion reveals differential regulation of cohesion establishment at the break site and genome-wide. Plos Genet 9:17.

39. Lau WCY, Li YY, Zhang QF, Huen MSY. 2015. Molecular architecture of the Ub-PCNA/pol eta complex bound to DNA. Sci Rep 5:15759.

40. Haracska L, Unk I, Prakash L, Prakash S. 2006. Ubiquitylation of yeast proliferating cell nuclear antigen and its implications for translesion DNA synthesis. Proc Natl Acad Sci USA 103:6477–6482.

41. Pabla R, Rozario D, Siede W. 2008. Regulation of Saccharomyces cerevisiae DNA polymerase eta transcript and protein. Radiat Environ Bioph 47:157–168.

42. Bienko M, Green CM, Sabbioneda S, Crosetto N, Matic I, Hibbert RG, Begovic T, Niimi A, Mann M, Lehmann AR, Dikic I. 2010. Regulation of translesion synthesis DNA polymerase eta by monoubiquitination. Mol Cell 37:396–407.

43. Alvarez V, Vinas L, Gallego-Sanchez A, Andres S, Sacristan MP, Bueno A. 2016. Orderly progression through S-phase requires dynamic ubiquitylation and deubiquitylation of PCNA. Sci Rep 6:25513

44. Bilsland E, Hult M, Bell SD, Sunnerhagen P, Downs JA. 2007. The Bre5/Ubp3 ubiquitin protease complex from budding yeast contributes to the cellular response to DNA damage. DNA Repair 6:1471–1484.

45. Mao P, Smerdon MJ. 2010. Yeast deubiquitinase Ubp3 interacts with the 26 S proteasome to facilitate Rad4 degradation. J Biol Chem 285:37542–37550.

46. Baudin A, Ozierkalogeropoulos O, Denouel A, Lacroute F, Cullin C. 1993. A simple and efficient method for direct gene deletion in Saccharomyces cerevisiae. Nucleic Acids Res 21:3329–3330.

47. Azevedo F, Marques F, Fokt H, Oliveira R, Johansson B. 2011. Measuring oxidative DNA damage and DNA repair using the yeast comet assay. Yeast 28:55–61.

48. Xu X, Lin A, Zhou C, Blackwell SR, Zhang Y, Wang Z, Feng Q, Guan R, Hanna MD, Chen Z, Xiao W. 2016. Involvement of budding yeast Rad5 in translesion DNA synthesis through physical interaction with Rev1. Nucleic Acids Res 44:5231–5245.

49. Tkach JM, Yimit A, Lee AY, Riffle M, Costanzo M, Jaschob D, Hendry JA, Ou J, Moffat J, Boone C, Davis TN, Nislow C, Brown GW. 2012. Dissecting DNA damage response pathways by analysing protein localization and abundance changes during DNA replication stress. Nat Cell Biol 14:966–976.

50. Fan Q, Xu X, Zhao X, Wang Q, Xiao W, Guo Y, Fu YV. 2018. Rad5 coordinates translesion DNA synthesis pathway by recognizing specific DNA structures in Saccharomyces cerevisiae. Curr Genet 64:889–899.

